# Inflated false negative rates undermine reproducibility in task-based fMRI

**DOI:** 10.1101/122788

**Authors:** G. Lohmann, J. Stelzer, K. Müller, E. Lacosse, T. Buschmann, V.J. Kumar, W. Grodd, K. Scheffler

## Abstract

Reproducibility is generally regarded as a hallmark of scientific validity. It can be undermined by two very different factors, namely inflated false positive rates or inflated false negative rates. Here we investigate the role of the second factor, i.e. the degree to which true effects are not detected reliably. The availability of large public databases and also supercomputing allows us to tackle this problem quantitatively. Specifically, we estimated the reproducibility in task-based fMRI data over different samples randomly drawn from a large cohort of subjects obtained from the Human Connectome Project. We use the full cohort as a standard of reference to approximate true positive effects, and compute the fraction of those effects that was detected reliably using standard software packages at various smaller sample sizes. We found that with standard sample sizes this fraction was less than 25 percent. We conclude that inflated false negative rates are a major factor that undermine reproducibility. We introduce a new statistical inference algorithm based on a novel test statistic and show that it improves reproducibility without inflating false positive rates.

## 1 Introduction

In recent years there has been a growing concern about the reliability and reproducibility of results obtained in human brain mapping using fMRI [1–5]. Two major factors contribute to this problem, namely inflated false positive rates [6] and lack of statistical power, i.e. inflated false negative rates [7–10]. While the detrimental effect of false positives is widely recognized, it may be less well known that inflated false negative rates may also diminish reproducibility, see Fig. 1 for an illustration.

**Figure 1:**
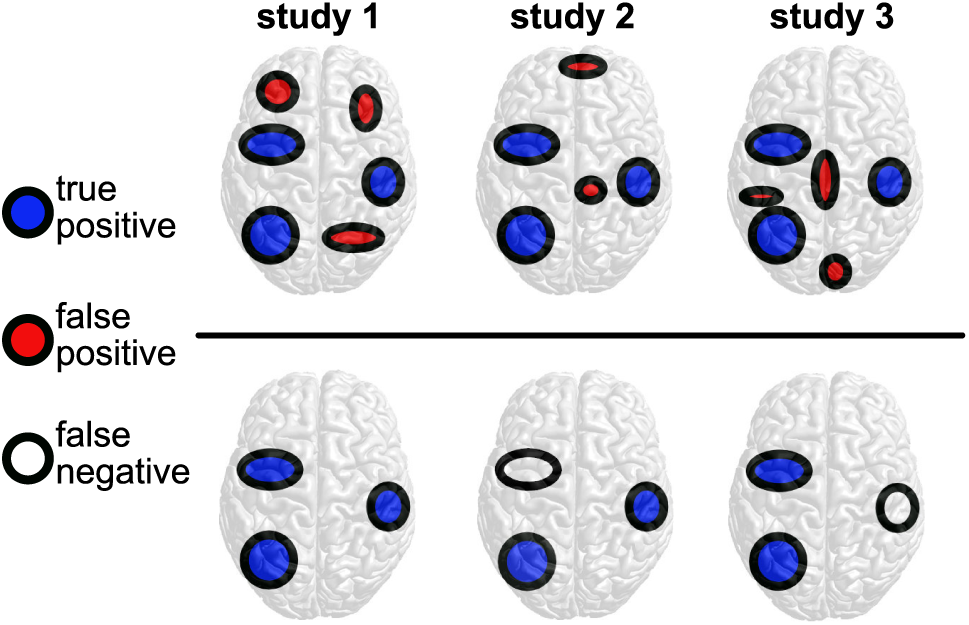
Illustration of the two factors that can undermine reproducibility across three repeats. Top row: false positives, bottom row: false negatives.

In the present context, we use the term *reproducibility* to denote robust detections over different samples drawn from one single large cohort. Specifically, we investigate the reproducibility in task-based fMRI data over different samples randomly drawn from a cohort of 400 subjects obtained from the Human Connectome Project (HCP) [11, 12]. Here we intentionally exclude factors such as different scanner hardware or different preprocessing regimes that might also diminish reproducibility, and focus instead on inter-subject variability and sensitivity of the test statistic as our main target of investigation.

Because of the absence of ground truth it is impossible to distinguish correct detections of true effects from erroneous detections. However, since high statistical power entails an increased likelihood that statistically significant results reflect true effects [7], it is reasonable to use the full cohort as a standard of reference for approximation. For brevity, we call it *GT400*. The GT400 map results from statistical inference corrected for the familywise error at *p* < 0.01, additionally thresholded to exclude very small effect sizes. The question we will address here is whether or not GT400 effects are robustly found at commonly used sample sizes. For this purpose, we will employ current methods as well as a new method that we introduce here. We will show that this new method improves sensitivity without inflating false positive rates.

We restrict ourselves to univariate activation maps in which activations can only be detected if their time courses correlate with a predefined hemodynamic response model [13, 14]. The multivariate and distributed nature of human brain function can however not be captured with these maps, so that an entirely new class of methods is now emerging [15–18]. Nonetheless, a large portion of our current neuroscientific knowledge relies on these classical results so that it is still relevant to check their validity.

A wide range of statistical inference methods have been proposed that aim to control error rates in fMRI-based human brain mapping, see e.g. [14, 19–23]. Eklund et al. [6] have argued that some of these methods may produce overly optimistic results, for a discussion see [24, 25]. Historically, the emphasis has been on controlling the false positive rate. In the present study, our focus will be on the false negative rate and the role it plays in diminishing reproducibility.

## 2 Results

We analysed task-based fMRI data provided by the Human Connectome Project (HCP), WU-Minn Consortium [11, 12]. We focused on two fMRI studies, namely the motor and the emotion task, using minimally preprocessed data of 400 participants. We generated an approximation to true positive effects as described in “Materials and Methods”. For ease of presentation, we call this approximation *GT400*. We estimated the percentage of GT400 that was actually found using three widely used statistical inference procedures FSL-TFCE [26], SPM-FWE, SPM-FDR [14, 27] and a new algorithm *LISA* that we introduce here. See “Materials and Methods” for more information.

### Approximate false negative rates using samples of size 20 (Figs. 2,3)

From a cohort of 400 subjects we randomly drew 100 samples of a size 20, and applied the four statistical inference procedures corrected for multiple comparisons at *p* < 0.05. We generated maps showing the reproducibility per voxel across these 100 tests (Fig. 2). Their histograms show the number of tests in which a given fraction of GT400 voxels was detected, see Fig. 3. We found that in the motor task, in 50 or more tests less than 20 percent of all GT400 voxels were detected. Thus, the approximate false negative rates were higher than 80 percent in more than half of the tests. In the emotion task, these rates were higher than 75 percent in more than half of the tests. With LISA, the detection rates and hence the estimated false negative rates were consistently better. In particular, LISA showed an increased detectability of the cerebellar (bilateral lobules HIV-VI and HVIII) and thalamic activation (right motor relay nuclei) during left-hand movement (motor task) as well as the improved thalamic and prefrontal and orbito-frontal activation in the emotion task. Note that SPM-FWE and SPM-FDR produced almost identical results. This may be due to the very stringent initial cluster forming thresholds *(CDT* = 0.001) [28].

**Figure 2:**
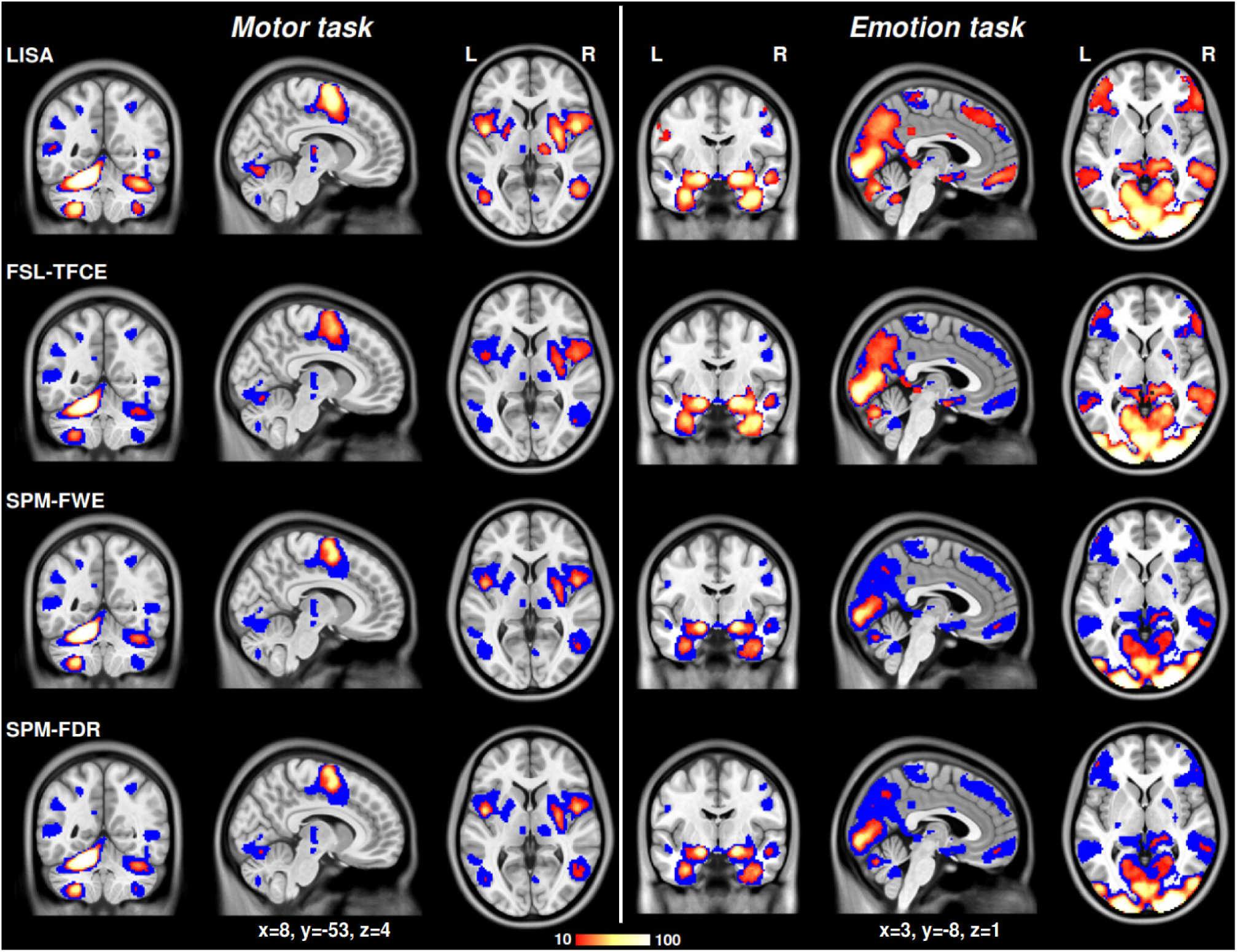
Reproducibility across 100 tests based on randomly drawn samples of size twenty. The colors represent the number of tests in which a voxel survived multiple comparison correction in at least 10 of the 100 tests. Thus, only areas in bright yellow or white are detected reliably, areas in dark red were only rarely detected. The underlying blue areas mark GT400 effects that were detected in less than 10% percent of all tests or not at all. We employed four different methods for statistical inference denoted as LISA, FSL-TFCE, SPM-FWE, and SPM-FDR, see "Materials and Methods”. Familywise error correction was applied for FSL-TFCE and SPM-FWE using p < 0.05 as threshold. Correction for the false discovery rate (FDR) was applied for LISA and also for SPM-FDR at p < 0.05. Note the increased detectability using LISA of the cerebellar (bilateral lobules HIV-VI and HVIII) and thalamic activation (right motor relay nuclei) during left-hand movement (motor task) as well as the improved thalamic and prefrontal and orbito-frontal activation in the emotion task. SPM-FWE and SPM-FDR yield almost identical results which may be due to the very stringent initial cluster-forming threshold (CDT=0.001). See also supplementary figure S3 for differences in reproducibility.

**Figure 3:**
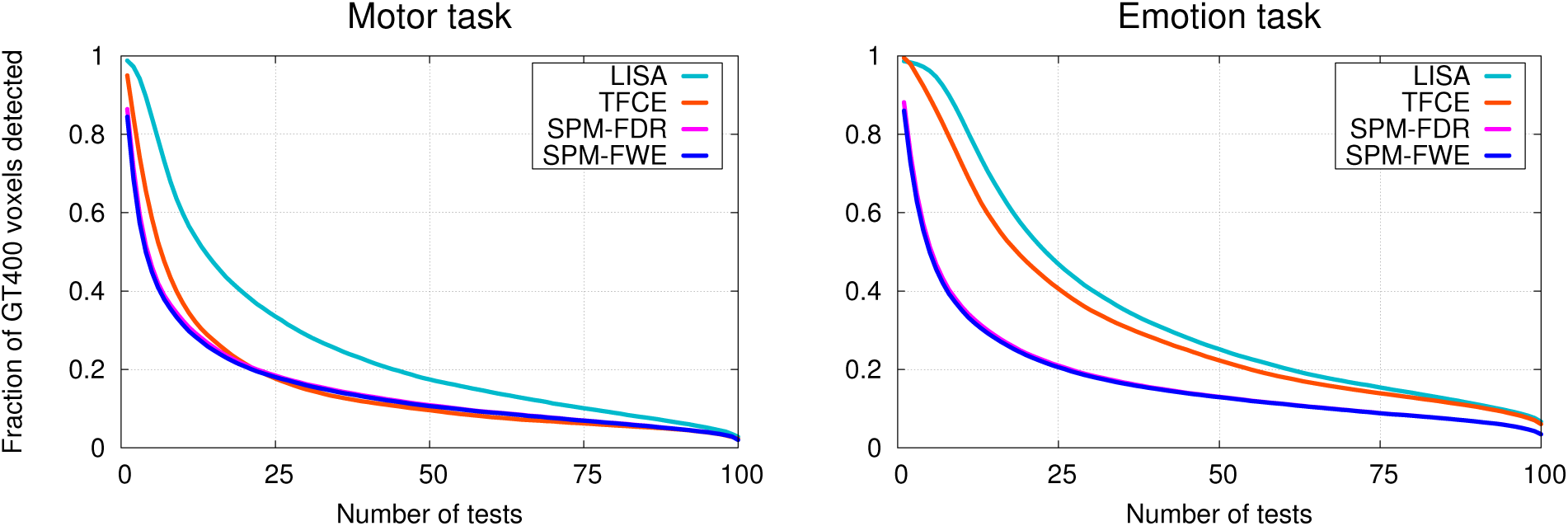
Approximate false negative rates at sample size 20. These histograms show the number of tests in which a given fraction of GT400 voxels was detected. For example, in the motor task, in 50 or more tests, less than 20 percent of all GT400 voxels were detected. Thus, in half of the tests the false negative rate is higher than 80 percent. In the emotion task, in half of the tests the false negative rates were higher than 75 percent. With LISA, these rates were consistently better.

### Rates of reliable detection across various sample sizes (Figs. 4,5)

We repeated the above reproducibility analysis for various samples sizes, namely 20, 40, 60, 80, and 100. We then recorded the sample size that was needed in order to detect an activation with reasonable chance of success which we defined to be 80 of the 100 tests (Fig. 4). We estimated detection rates as follows. We counted the number of voxels in each of the sample size maps of Fig. 4 as well in the GT400 map and weighted each voxel by its effect size so that voxels with low effects contribute less to the final result. We define the “effectsize weighted detection rate” as the ratio of these weighted counts. Effect sizes are defined as the voxelwise mean effect across all 400 subjects divided by the standard deviation, see also “Materials and Methods”. Figure 5 shows that with sample sizes of 20, less than 25 percent of the total GT400 effect is reliably detected. Even with sample sizes of 60, detection rates are below 55%, see also supplementary table S2. Note that generally smaller sample sizes are needed using the algorithm LISA, see supplementary figure S4.

**Figure 4:**
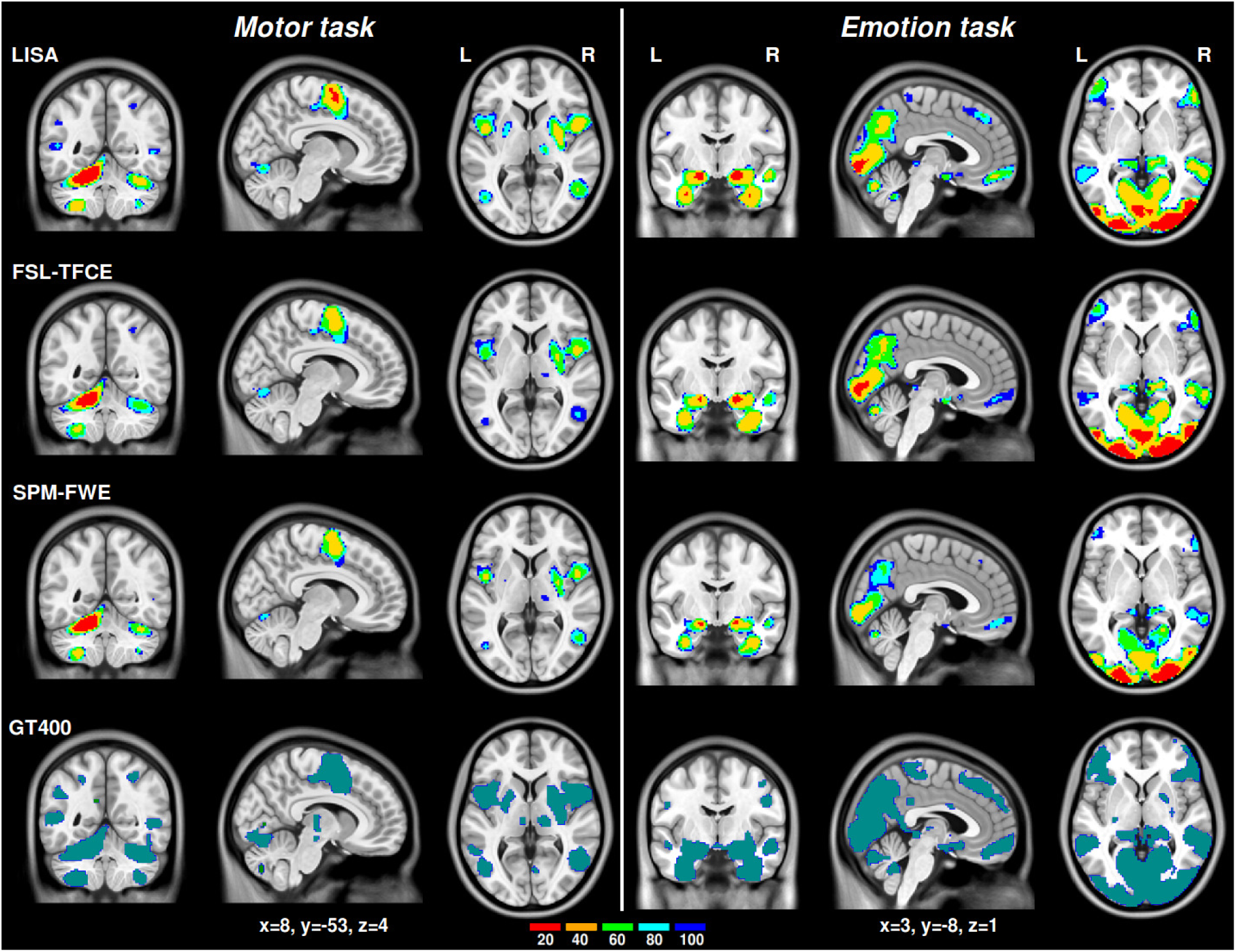
Maps indicating minimal sample sizes required for reliable detection. The colors indicate the minimal sample sizes that are needed in order to detect an activation with reasonable chance of success, i.e., in 80 of 100 tests. The results for SPM-FDR were almost identical to those of SPM-FWE. The GT400 map is shown in the bottom row.

**Figure 5:**
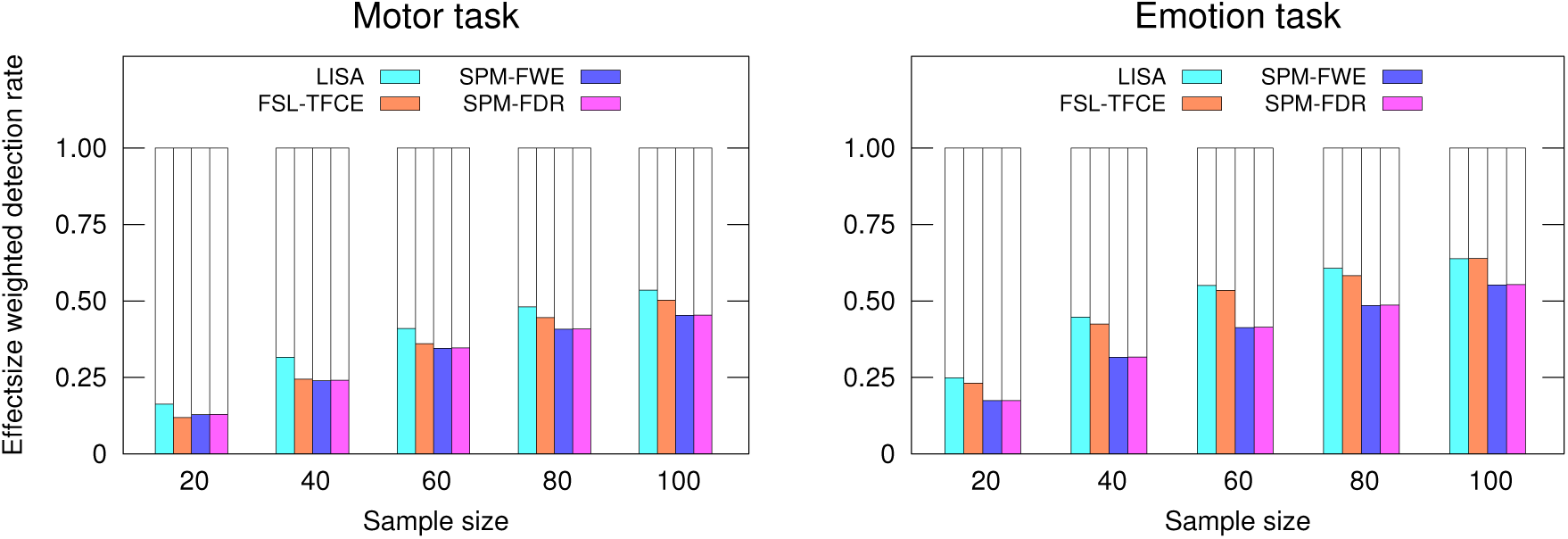
Effectsize weighted detection rates. The histograms show the fraction of the overall effect within GT400 that was reliably detected, i.e. in at least 80 of 100 tests. Each voxel was weighted by its effect size, so that voxels with low effects contribute less to the total sum. At sample sizes of 20, less than 25 percent of the total GT400 effect was detected.

## 3 Conclusions

In this study we derived approximations for the false negative rates in two HCP studies using standard procedures as well as the new method *LISA*. The most important finding is that with commonly used sample sizes of 20 the false negative rates exceeded 75 percent in more than half of 100 tests (Fig. 3). Furthermore, less than 25 percent of the overall GT400 effect was detected reliably (Fig. 5). With larger sample sizes the detection rates improve but are still far away from the costumary 80% level as a standard of adequacy for the type II error. We used an effectsize weighted measure for the detection rate so that type II errors that may be considered less relevant, e.g. around the borders of activation areas, do not distort the results.

These results lead us to conclude that inflated false negative rates do indeed undermine reproducibility. The recent publication by Eklund et al. [6] has drawn a lot of attention to the problem of inflated false positives. However, because of the large sizes of the false negative rates, we hypothesize that false negatives may actually play a much more important role in diminishing reproducibility. If true, this would be quite reassuring because it entails that most nonreproducible results may in fact not be false, but simply based on effects that are too weak to be detected reliably.

In a recent study, Poldrack et al. [1] reported that median sample sizes have steadily increased since 1995, but even in 2015 they were below 30. Our findings show that even larger samples are too small for a reliable detection of all true effects. Data acquisition of large samples is time consuming and costly, and may often become impracticable. Therefore, improvements in reproducibility cannot be achieved by increasing sample sizes alone. Rather, new analysis methods with improved sensitivity and reliability are crucial.

Here we have introduced the new method *LISA* which is based on a novel test statistic derived from hotspot analysis in geographical information systems. In our experiments, we found that LISA was more sensitive compared to the other three methods and required considerably smaller samples to achieve similar results. Importantly, LISA passed the test proposed by Eklund et al. [6] so that we may assume that the decrease in false negatives did not lead to an inflation in false positives. However, even though LISA outperformed the other three methods in terms of sensitivity, it was still far away from uncovering all GT400 effects.

In LISA, hotspots of activation are highlighted as locally coherent regions. Local coherence as a new feature for statistical inference is very attractive because it is model free and easily accessible with neuroimaging techniques. More importantly, its effectiveness may point to an underlying very general mechanism of brain function. We have previously proposed a similar feature of local coherence for network detection where it also proved to be a very strong marker of differential task involvement [18].

The many small sized samples drawn from the total cohort of 400 subjects are likely to have overlapped considerably. In such a scenario one would expect a high degree of reproducibility that is simply due to the correlation across samples. In more realistic settings where such overlap is absent, and other factors such as different scanner hardware or different preprocessing regimes play a role, reproducibility may be even worse. Therefore, our results are more likely to err on the conservative side. We should note however that the degree of overlap varies with sample size, so that it is not possible to directly compare reproducibility values across different sample sizes. But with larger sample sizes and hence larger overlap across samples the degree to which we underestimate the problem is likely to increase.

Based on the results reported by Eklund et al. [6], we used very stringent initial cluster forming thresholds for SPM-FWE and SPM-FDR (CDT=0.001). A first consequence of this choice was that both methods produced very similar results [28]. More importantly however, this choice has lead to a dramatic increase in false negatives and hence to a loss of reproducibility. Thus, an overemphasis on avoiding false positives can be problematic.

In summary, we conclude that inflated false negative rates are a major impediment for reproducible human brain mapping. Since inflated false negative rates may lead to an incomplete and hence biased picture of brain function, it should be seen as a problem that is just as severe as inflated false positive rates.

## 4 Materials and Methods

### HCP data acquisition

All data sets were acquired with the following parameters: TR=720ms, TE=33.1ms, 2 mm isotropic voxel size, multiband factor 8. We focused on task-based fMRI tasks, namely the motor and the emotion task, using minimally pre-processed data of 400 participants of the left-right phase-encoding runs. The preprocessing protocol is described in [29]. In addition, FSL-Fix [30–32] was used for data denoising, removing physiological nuisance effects as well as motion and multi-band artifacts. In addition, we applied a Gaussian spatial filter with a kernel size of 6 mm. Using FSL-FEAT [30–32], we performed individual onesample t-tests resulting in 400 single-subject maps of uncorrected z-values.

### HCP Motor task

While in the scanner, participants were cued visually to tap their left or right fingers, squeeze their left or right toes, or move their tongue. Each block lasted 12 seconds (10 movements), and was preceded by a 3 second cue, for details see [12]. Here we investigated only the left hand fingertapping condition.

### HCP Emotion task

In this experiment, participants were cued to decide which of two faces matched the face shown on top of the screen. In a second experimental condition, an analogous task was done using shapes instead of faces. The faces had either angry or fearful expressions, for details see [12]. Here we investigated the contrast “faces minus shapes”.

### Existing statistical inference procedures

Cluster-extent based thresholding schemes are currently the most popular methods for statistical inference in fMRI [33]. Therefore, we selected a commonly used method of this type for reference, namely the Gaussian Random Field method as implemented in SPM12 [14,27,34], with corrections for either the familywise error (SPM-FWE) or for the false discovery rate (SPM-FDR). In our experiments, we used the default initial cluster forming threshold of CDT=0.001. With a lower threshold, the test reported by Eklund et al. [6] produced inflated false positive rates. As a representative of a completely different method, we choose threshold-free cluster enhancement (FSL-TFCE) implemented in FSL [26, 31]. Even though many other methods exist, we restricted ourselves to the ones listed above because they represent a large portion of the existing literature.

### Effect sizes

Since large sample sizes may produce highly significant results with very low effect sizes, we computed the voxelwise effect sizes defined as the mean across all 400 data sets divided by their standard deviation. Following Cohen [35], effect sizes of 0.2 are regarded as small, 0.5 as medium and 0.8 as large.

### A new statistical inference procedure *LISA*

We introduce a new method of second-level statistical inference for task-based fMRI called *LISA*. The input into LISA is a set of several single-subject maps. The output is a single thresholded map representing a voxel-wise onesample t-test controlled for a predefined false discovery rate. As a first step, LISA performs a one-sample voxelwise t-test across all input maps in which each voxel receives a t-value uncorrected for multiple comparisons. In a second step, a bilateral filter is applied to this map [36, 37]. The effect of this filter is to highlight hotspots of activation defined as areas of local concentrations of large t-values. The bilateral filter replaces each voxel with a weighted average of similar and nearby voxel values. It uses a product of two Gaussian smoothing kernels where one kernel penalizes spatial distance while the other penalizes discrepancies in voxel values. Thus, a voxel only receives a high weight if it is both spatially close to the center of its local neighbourhood and its value is close to that of the center voxel. The bilateral filter is defined as follows. Let *i* denote some voxel position, Ω its local neighbourhood, and *z_i_* its value in the map of uncorrected z-values. Then the filtered value *λ_i_* is given by

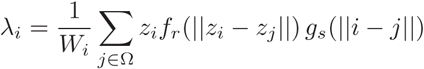

with normalizing factor

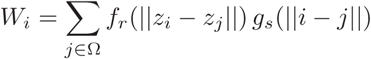

where *f_r_* is the range kernel for voxel intensities, and *g_s_* is a spatial kernel for for weighting differences in voxel coordinates. As customary, we use Gaussian kernels for *f_r_* and *g_s_* with *f_r_*(*x*) = *exp*(*−x^2^/σ_r_*) and *g_s_*(*x*) = *exp*(*−x^2^*/*σ_s_*). Note that this filter requires three parameters, namely the size of the local neighbourhood Ω, and *σ*_*r*_, *σ*_*s*_ for the kernel functions *f_r_, g_s_*. We determined values for these parameters using simulated data. Once determined, these settings were kept constant in all our experiments, regardless of spatial resolution or smoothing parameters used during preprocessing. Specifically, we used *σ_s_* = 2.0 and a neighbourhood shaped like sphere with a radius of two voxels comprising 57 voxels. The parameter *σ_r_*, must however be scaled by the global variance *varglobal* of the map of uncorrected z-values. We then use *σ_r_* = 1.9 × *var_global_* throughout. Bilateral filtering can be applied iteratively. In all our experiments, we used two iterations. The filtered values *λ_i_* are used as a test statistic for statistical inference.

LISA controls the false discovery rate (FDR) using random permutations. In onesample tests, signs are reversed in randomly selected maps. The false discovery rate is then estimated using a two-component model of the form *F dr* = *p_0_F_0_/F_z_* where *F*_0_ is the empirical null distribution derived from random permutations, *F_z_* is the distribution of voxel values after bilateral filtering of the non-permuted map, and *p*_0_ is the prior probability of the null [38]. For simplicity, we use *p*_0_ = 1 which is the most conservative choice. In the experiments reported here, we used 5000 permutations. A more detailed description of this algorithm can be found in (t.b.a)˙.

### The Eklund test applied to LISA

To make sure that LISA does not produce inflated false positive rates, we subjected it to the test described by Eklund et al. [6]. We found that error rates were well within the target range of *p* < 0.05. For details see the supplementary material.

### Approximation to ground truth (GT400)

Since ground truth is not available, it is not possible to directly assess the false negative rate. However, a reasonable approximation can be obtained as follows. We use FSL-TFCE as a reference method because it is widely used and validated [26] and apply statistical inference using the full cohort of 400 subjects as a sample. Note that high statistical power entails an increased likelihood that statistically significant results reflect true effects [7]. We correct for the familywise error at *p* < 0.01 using 10000 random permutations so that the probability of type I errors is less than one percent. To avoid statistically significant but very small effects, we discard voxels with effect sizes below 0.2. For brevity, we will call this approximation *GT400*. The resulting GT400 maps are shown in supplementary figures S1,S2.

### Effect-size weighted detection rates

We computed weighted sums of the voxels in each of the sample size maps of figure 4 and in the GT400 map. Each voxel was weighted by its effect size, so that voxels with low effects contribute less to the total sum. We found that with sample sizes of 20, less than 25% of such effects were reliably detected (Fig. 5).

### Software

The software for the LISA algorithm is available at (t.b.a.).

## Acknowledgments

This work was partly funded by the European Commission H2020-PHC-2014 634541 CDS-QUAMRI.

Data were provided by the Human Connectome Project, WU-Minn Consortium (Principal Investigators: David Van Essen and Kamil Ugurbil; 1U54MH091657) funded by the 16 NIH Institutes and Centers that support the NIH Blueprint for Neuroscience Research; and by the McDonnell Center for Systems Neuroscience at Washington University.

Computations were partly performed at the Max Planck Computing and Data Facility, Garching, Germany.

## Supplementary Information

### The Eklund test applied to LISA

To make sure that LISA does not produce inflated false positive rates, we subjected it to the test described by Eklund et al. [6]. We used the same experimental designs (B1,B2,E1,E2) and preprocessing regimes, and applied LISA to all 200 data sets of the Beijing sample of [6]. The preprocessing pipeline is based on SPM12 [34]. For each of the four experimental designs and four spatial smoothing kernels (4mm, 6mm, 8mm 10mm), we randomly drew 1000 samples consisting of 20 data sets each and applied LISA as described above. To allow for a direct comparison with the results of [6], we recorded the family wise error rate. To check whether LISA is vulnerable to resampling parameters used during preprocessing, we repeated the entire Eklund test with the resampling parameter set to (*3mm*)^3^ instead of (*2mm*)^3^ while parameter settings for the bilateral filter remained identical. In both cases, we found that error rates were well within the target range of *p* < 0.05. The results are shown in table S1.

**Table S1:** Results of the Eklund test using LISA. The table shows false positive rates of the Eklund test with two types of preprocessing where the resampling parameter was either set to (2 mm)^3^ or to (3mm)^3^ resolution. Both results are based on 4x4x1000 individual tests. The experimental designs (E1,E2,B1,B2) are the same as in Eklund et al. [6]. For preprocessing, SPM12 was used. Four levels of spatial smoothing were used during preprocessing (4mm, 6mm, 8mm, 10mm).

| | $(2mm)^3$ resolution | | | | $(3mm)^3$ resolution | | | |
| --- | --- | --- | --- | --- | --- | --- | --- | --- |
|  | E1 | E2 | B1 | B2 | E1 | E2 | B1 | B2 |
| 4mm | 0.007 | 0.001 | 0.020 | 0.019 | 0.000 | 0.003 | 0.022 | 0.017 |
| 6mm | 0.020 | 0.006 | 0.037 | 0.027 | 0.011 | 0.009 | 0.047 | 0.017 |
| 8mm | 0.026 | 0.013 | 0.060 | 0.033 | 0.029 | 0.007 | 0.049 | 0.035 |
| 10mm | 0.037 | 0.017 | 0.054 | 0.035 | 0.032 | 0.014 | 0.065 | 0.030 |

**Figure S1:**
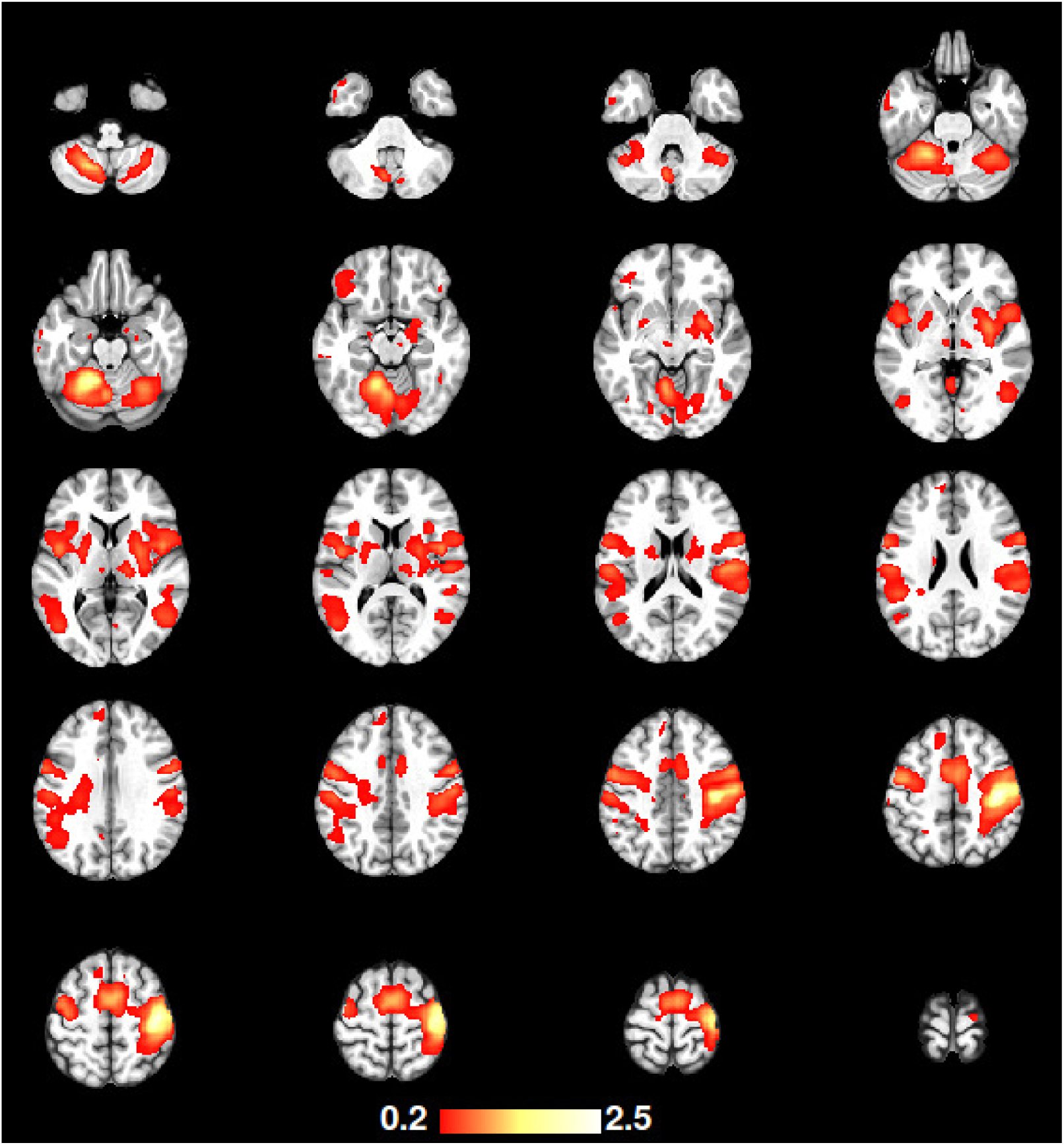
The GT400 map of the motor task. The GT400 map shows voxels that are detected at p < 0.01 FWE-corrected using FSL-TFCE using a sample size of 400. This map is thresholded using effect sizes larger than 0.2 where effectsize is defined as the voxelwise mean across all 400 subjects divided by the standard deviation. The colors encode effectsize.

**Figure S2:**
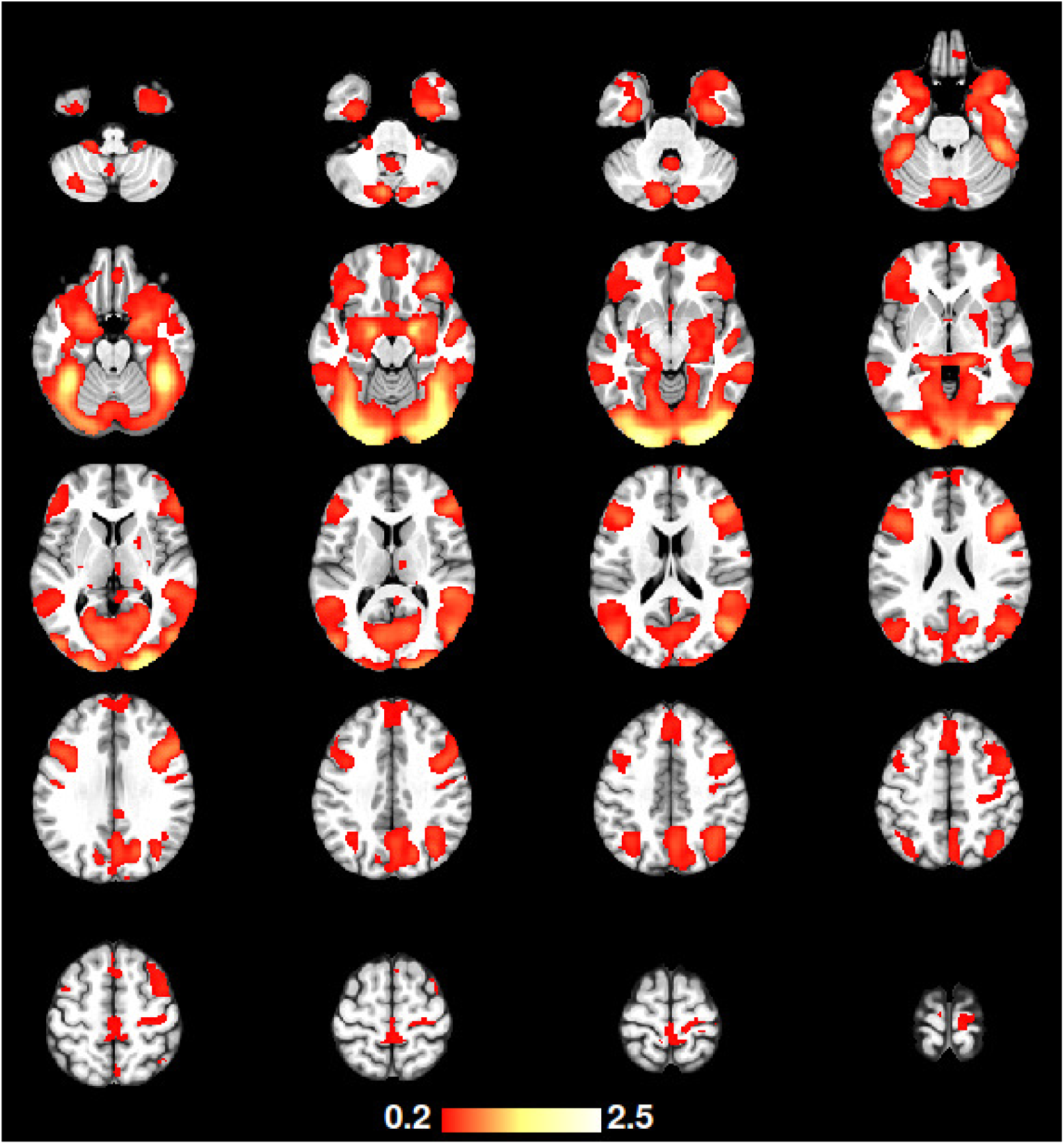
The GT400 map of the emotion task. The GT400 map shows voxels that are detected at p < 0.01 FWE-corrected using FSL-TFCE using a sample size of 400. This map is thresholded using effect sizes larger than 0.2 where effectsize is defined as the voxelwise mean across all 400 subjects divided by the standard deviation. The colors encode effectsize.

**Table S2:**
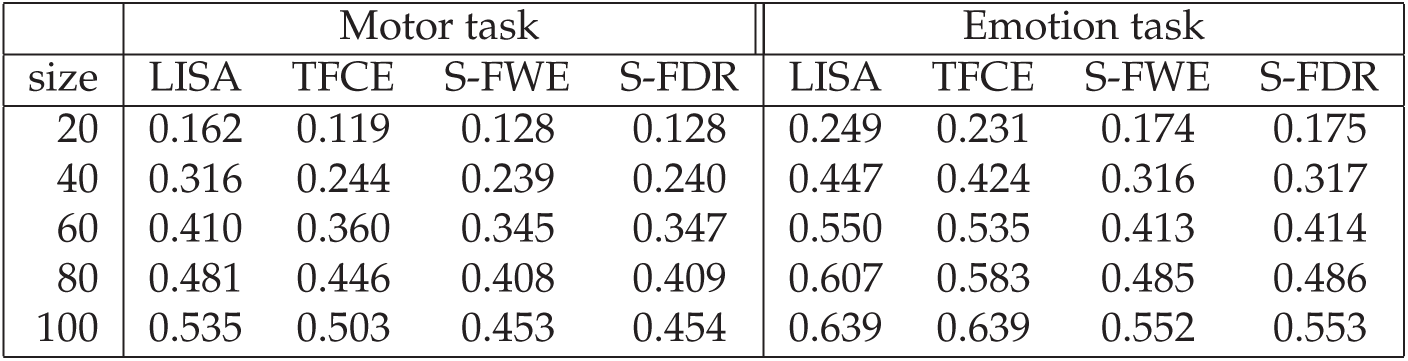
Effectsize weighted detection rates. This table presents the same data as figure 5.

|  | Motor task |  |  |  | Emotion task |  |  |  |
| --- | --- | --- | --- | --- | --- | --- | --- | --- |
| size | LISA | TFCE | S-FWE | S-FDR | LISA | TFCE | S-FWE | S-FDR |
| 20 | 0.162 | 0.119 | 0.128 | 0.128 | 0.249 | 0.231 | 0.174 | 0.175 |
| 40 | 0.316 | 0.244 | 0.239 | 0.240 | 0.447 | 0.424 | 0.316 | 0.317 |
| 60 | 0.410 | 0.360 | 0.345 | 0.347 | 0.550 | 0.535 | 0.413 | 0.414 |
| 80 | 0.481 | 0.446 | 0.408 | 0.409 | 0.607 | 0.583 | 0.485 | 0.486 |
| 100 | 0.535 | 0.503 | 0.453 | 0.454 | 0.639 | 0.639 | 0.552 | 0.553 |

**Figure S3:**
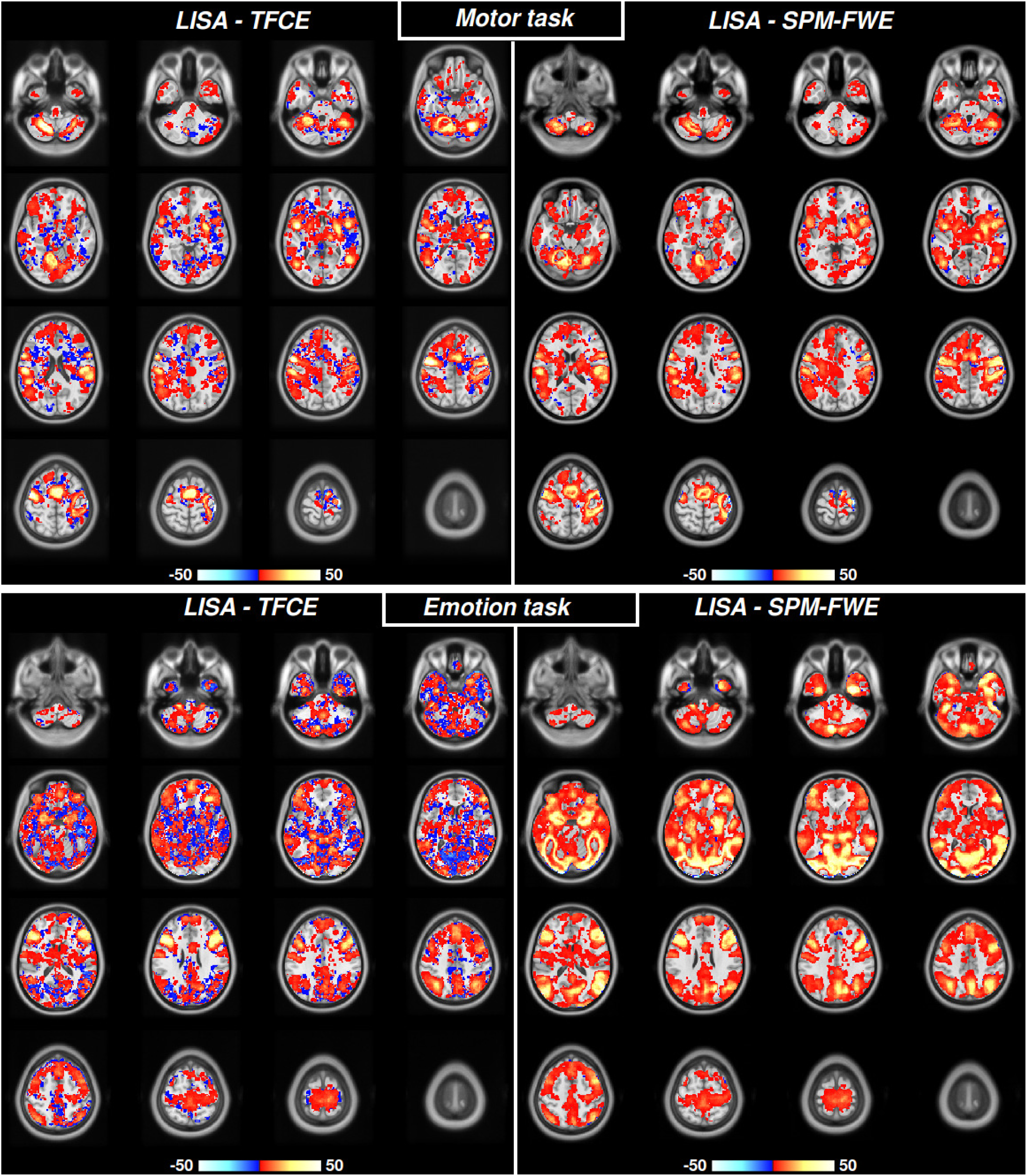
Difference of reproducibility maps of figure 2. The comparison with SPM-FWE shows that LISA has better reproducibility almost everywhere, indicated by the positive (red-yellow) color code. The comparison with SPM-FWE shows that LISA has better reproducibility particularly in GT400 activation areas.

**Figure S4:**
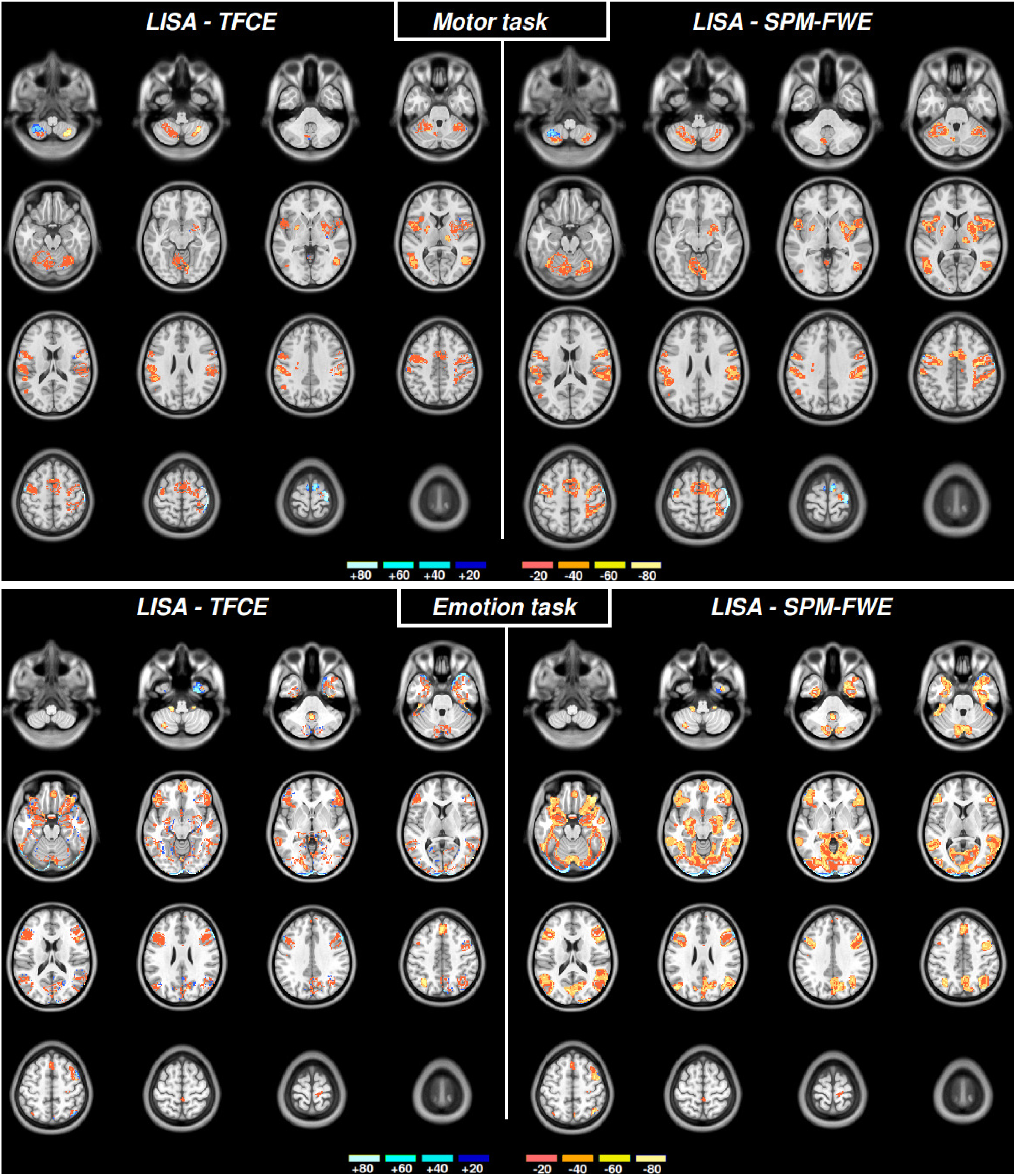
Difference LISA-TFCE of sample size maps of figure 4. The comparison with SPM-FWE and FSL-TFCE shows that LISA needs smaller sample sizes almost everywhere, indicated by the positive (red-yellow) color code.

